# Strategies used by two memories to share space in a common neural network

**DOI:** 10.1101/2025.04.07.647298

**Authors:** Viral K. Mistry, Evan S. Hill, William N. Frost

**Affiliations:** Stanson-Toshok Center for Brain Function and Repair, Rosalind Franklin University of Medicine and Science, North Chicago, IL, 60064; School of Graduate and Postdoctoral Studies, Rosalind Franklin University of Medicine and Science, North Chicago, IL, 60064; Chicago Medical School, Rosalind Franklin University of Medicine and Science, North Chicago, IL, 60064

## Abstract

How distinct memories are encoded into the same network space without destructive interference is not well-understood. Here, we utilized *Tritonia diomedea*’s escape swim network to explore how two sequentially acquired forms of non-associative learning, sensitization and habituation, are encoded into the same network. Behavioral experiments showed them to alter non-identical sets of behavioral features, suggesting they utilize somewhat independent sites of plasticity within the network. Large-scale voltage-sensitive dye recordings revealed two findings. First, both forms of learning, which occur sequentially in the 10-trial training protocol used, act to produce a change in the number of pedal neurons firing during the dorsal phase of the motor program, with sensitization producing an increase, and habituation a decrease in their number. The number of neurons participating in the ventral phase was unaffected. Second, sensitization produced an enhancement of burst intensity specific to the ventral phase neurons, while habituation was associated with a decrease in burst intensity in both phases. Using injected current pulses, intracellular recordings revealed that sensitization acted to increase the excitability of neurons firing in both phases, whereas habituation only acted to reduce excitability in ventral phase neurons. These excitability changes were associated with different mechanisms – reduced spike frequency accommodation in the ventral phase neurons, and depolarization of the resting potential in the dorsal phase neurons. These findings of partially different storage sites and mechanisms for two different non-associative memories illuminate a potential network strategy for minimizing destructive interference when storing multiple memories into the same network.

## INTRODUCTION

When learning occurs, neural networks alter their properties to encode the new information into memory traces, which constitute the physical substrate of learning [1]. Investigating how memory traces are formed and maintained has been greatly facilitated by technical advances that enable precise interrogation of the specific alterations within neural networks that underpin memory formation. Such changes include modifications to intrinsic excitability, synaptic strength and the numbers of participating neurons [2, 3]. A key question arises when attempting to generalize these findings beyond a single memory – how can multiple memories be successfully encoded into the same network without interfering with one another? Work in overlapping memories in rodent associative learning paradigms suggest that in order for a network to keep different memories distinct, they must utilize distinct neurons and synapses to preserve their fidelity and specificity [4–6]. However, this work has relied on indirect measures of neuronal activity, such as calcium activity or immediate early gene expression, as the complexity and scale of mammalian neural networks makes direct observation challenging. As a result, significant gaps remain in our understanding of the specific information encoded by these changes and the mechanisms through which this encoding occurs [7].

Marine invertebrates have a well-established history as model organisms for studying the cellular and molecular basis of learning [8, 9]. While some of this work has focused on associative learning paradigms [10, 11], much of it has examined non-associative learning, the most fundamental form of learning observed in animals [12–14]. Two key types of non-associative learning are sensitization, where a novel aversive stimulus heightens behavioral responses, and habituation, where repeated exposure to the same stimulus diminishes the behavioral response. Studies of how these forms of non-associative learning are encoded in the neural networks of simple marine invertebrates have uncovered evolutionarily conserved principles of memory formation, from changes to the properties of individual neurons to the reconfiguration of entire neural networks [15–17].

In this study, we leverage the nudibranch mollusk *Tritonia diomedea* and its escape swim motor program (SMP) to examine for the first time how two sequential forms of non-associative learning are encoded into the same escape network. When *Tritonia* is attacked by its predator, *Pycnopodia helianthoides*, it initiates an escape swim consisting of rhythmic ventral-dorsal full body flexions that propel it away to safety (**Fig. 1A**) [18]. This behavior can be readily elicited in the laboratory by applying concentrated salt to the animal’s tail [19]. *Tritonia*’s SMP network has been well characterized: sensory input activates a command neuron that in turn excites a central pattern generator (CPG) that produces motor output via a large population of flexion neurons [20]. The escape swim has multiple quantifiable behavioral parameters and readily undergoes non-associative learning in both intact animals and in isolated brain preparations [21, 22]. Sensitization learning in *Tritonia*’s SMP has been previously shown to shorten swim onset latency, and increase both cycle number and the animal’s initial jump height off the substrate as the swim commences, while habituation has been shown to lengthen response latency and decrease cycle number [23–26]. Both forms of learning are relevant to the animal – sensitization improves the behavioral response in order to avoid predation if a follow-up attack occurs, while habituation weakens the response to avoid wasting energy on unimportant stimuli [25]. Importantly, these two forms of learning have been shown to coexist – swim onset latency can remain sensitized even as cycle number has decreased, demonstrating that these two memories can occupy the same network without destructive interference [24]. This potential for coexisting memories provided the opportunity to explore a fundamental problem for all brains: how multiple memories can be stored in common networks while minimizing destructive interference.

**Figure 1.**
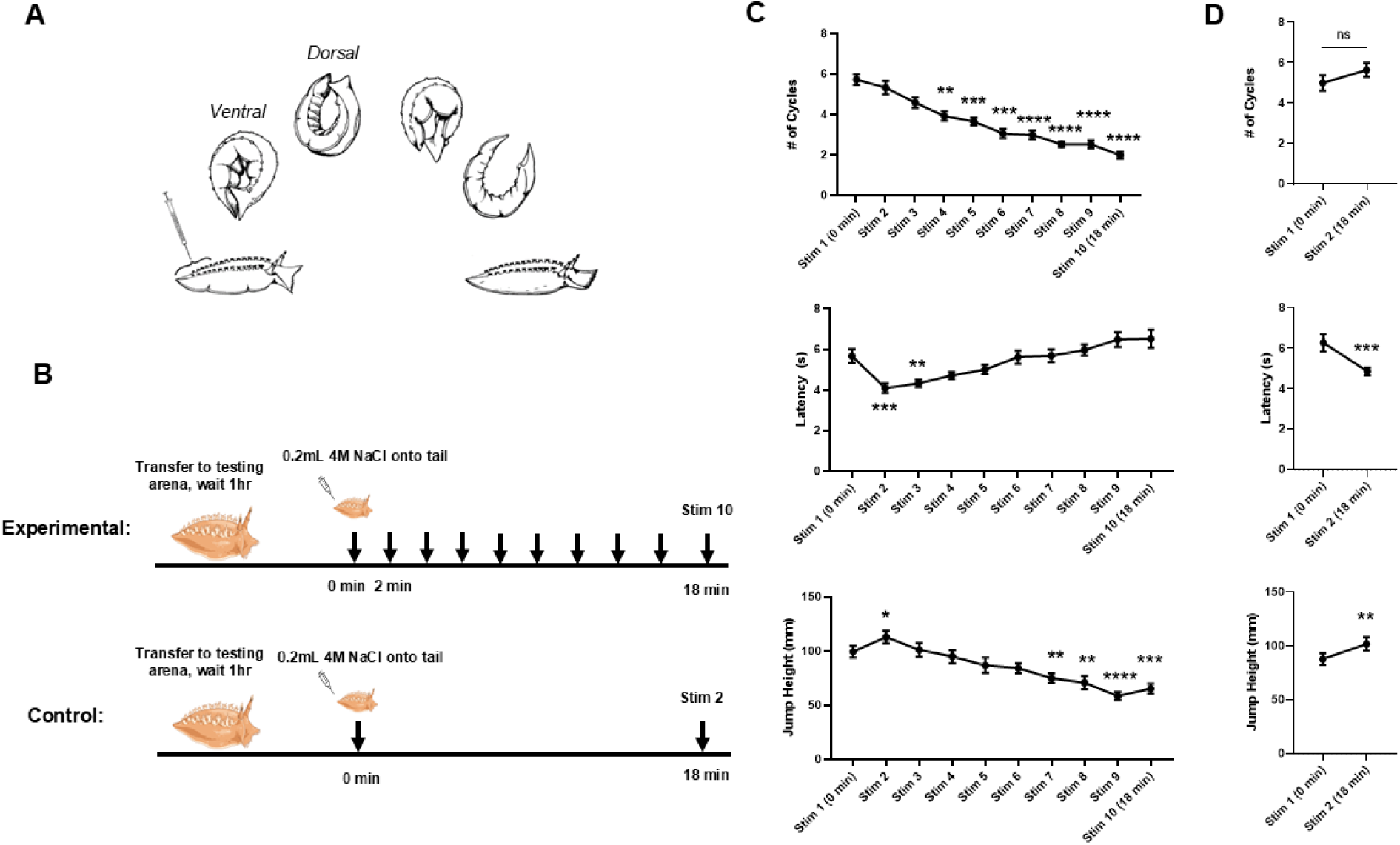
Behavioral parameters of the escape swim differentially undergo non-associative learning. **A)** Illustration depicting the stages of a *Tritonia* escape swim initiated via concentrated salt stimulus to the tail, resulting in an initial ventral flexion followed by a dorsal flexion, that repeats until the animal returns to the substrate. **B)** Protocol for behavioral learning (image of *Tritonia* from BioRender). **C)** Jump height, onset latency, and cycle number each differentially change at different time points with repeated stimuli within the experimental group (n = 15). ** indicates p < 0.01, *** indicates p <0.001, **** indicates p < 0.0001. **D)** Jump height, onset latency, and cycle number in the control condition (n = 17). Onset latency and jump height sensitization persist for at least 18 minutes in intact animals, while cycle number remains unchanged. ** indicates p < 0.01, *** indicates p <0.001. Error bars represent SEM.

We have demonstrated here how *Tritonia*’s escape network employs distributed sites of plasticity in the network to encode sequential memories for sensitization and habituation. This includes modifications to the number of participants, flexion burst intensity, excitability, and resting membrane potential. These changes differed in both their mechanisms and flexion phase, representing unique strategies for encoding plasticity. While the two memories did appear to directly conflict at some sites, resulting in some destructive interference, their use of independent sites and their employment of different mechanisms when overlapping in the same neurons provides a framework of how multiple memories can be encoded into the same limited network space while minimizing interference.

## METHODS

### Preparation

*Tritonia diomedea* were obtained from Living Elements LLC (Vancouver, CA). Animals were maintained in chilled (9.5-11.5°C) recirculating artificial seawater systems prior to experiments. The artificial seawater (ASW) was made using Instant Ocean mix. Experiments involving isolated brains were performed in filtered ASW.

### Behavioral experiments

*Tritonia* were weighed and then placed alone into the top tier of a plexiglass recirculating seawater tank. 1 hour after being moved into the testing region, swims were elicited by applying 0.2mL of 4M NaCl to the tail of the animal. Escape swims were filmed with a digital camcorder. A ruler was placed near the animal in the tank to allow measurement of the height of the initial jump off the substrate powered by the first ventral flexion of the swim. Afterwards, videos were analyzed using the same method as [26]: the height of the jump was measured as the maximal height achieved by the center of the animal on the first cycle, and the onset latency was measured as the time between the application of the NaCl solution to the maximum point of the first ventral flexion.

### Optical recordings

Simultaneous action potential activity in large numbers of neurons were recorded using a voltage sensitive dye and photodiode array, as in [26–28]. *Tritonia*’s central ganglia, consisting of the bilateral cerebropleural and pedal ganglia were dissected and pinned to the bottom of a Sylgard (Dow Corning) lined Petri dish containing filtered ASW. During the entire experiment, the temperature was maintained between 9.0 and 10.0°C by passing filtered ASW through a feedback-controlled in-line Peltier cooling system (Model SC-20, Warner Instruments) using a peristaltic pump (Model 720, Instech Laboratories). Temperature was monitored with a BAT-12 thermometer fitted with an IT-18 microprobe (Physiotemp Instruments). The central ganglia were then partially fixed in 0.25% glutaraldehyde for 10 seconds to limit prep movement. Pedal ganglion neurons were stained by periodically applying pressure to a PE tube filled with 0.2mg/mL RH-155 (Toronto Research Corporation, Toronto, CA), pressed against the surface of the ganglion for 1 hour. Following staining, the pedal ganglion was transferred and pinned to a Sylgard-lined perfusion chamber used for optical recording, where the PdN3 contralateral to the stained pedal ganglion was pulled into a suction electrode. To improve visualization, two small pieces of silicone were placed in the chamber on opposite sides of the stained ganglion, allowing for a glass coverslip fragment to be pressed down and held in place to partially flatten the stained pedal ganglion (as described and visualized in [27]). The preparation was trans-illuminated by a 735nm LED (Thor Labs), and the light was collected by a 20x 0.95 NA water-immersion objective lens (Olympus) and passed through a phototube to reach either the photodiode array or a camera (Optronics). The preparation rested on the optical imaging rig for 90 minutes following nerve suction before acquisitions began. Fictive swim motor programs were elicited via stimulation of the contralateral PdN3 (7V, 1Hz, 7s stimulus train, 5ms pulses). In the experimental group, 10 stimuli were delivered with 2-minute inter-stimulus intervals. In the control group, two stimuli were delivered with an 18-minute inter-stimulus interval. Voltage activity was captured using a RedShirtImaging PDA-III photodiode array consisting of 464 photodiodes sampled at 1600 Hz. Data were acquired using RedShirtImaging’s Neuroplex software. Optical data were AC-coupled and then amplified 100x by the PDA-III. After acquisition, optical data were band-passed filtered in the Neuroplex software (Butterworth, 5Hz high pass, 100Hz low pass), and saved as text files. Independent component analysis (ICA) was performed on the filtered optical data in MATLAB to yield a set of single neuron action potential traces, as described in [29]. Following ICA, the resulting components had a manual threshold applied to generate binary records of action potential spike times for each neuron.

### Analysis of spike trains from optical recordings

Bursters were determined using the method described in [26, 30]: An algorithm identified all neurons that fired at least one burst during the SMP using power spectral density. Neurons that fired on the first cycle of every swim motor program were then used for calculating burst intensity. The spike time files were analyzed using a custom MATLAB script to identify the number of spikes in the first burst of each bursting neuron. Candidate bursters from each phase were used to identify the start and end of the first flexion. Ventral and dorsal phase were identified relative to the stimulus – dorsal phase neurons can tonically fire with the incoming stimulus, but the first complete burst is the ventral phase, followed by the dorsal phase.

### Intracellular recordings

Intracellular electrophysiology recordings were obtained from separate, non-stained isolated brain preparations with 10-25 mΩ electrodes filled with 3M KCl. Electrodes were positioned with Sensapex micromanipulators and connected to Dagan IX2-700 intracellular amplifiers. During the entire experiment, the temperature was maintained between 9.0 and 10.0°C by passing filtered ASW through a feedback-controlled in-line Peltier cooling system (Model SC-20, Warner Instruments) using a gravity perfusion bottle. Temperature was monitored with a BAT-12 thermometer fitted with an IT-18 microprobe (Physiotemp Instruments). Neuronal recordings were digitized at 2 KHz with a BIOPAC MP150 data acquisition system in Acqknowledge 3.9.1. Swim motor programs were elicited by stimulation of the contralateral PdN3 relative to the pedal ganglion that was desheathed (7V, 1Hz, 7s stimulus train, 5ms pulses). Once a stable recording was obtained, intrinsic excitability was determined by injecting three 5s depolarizing constant current pulses into the cell, with 5s intervals between current injections, using an AMPI Master-8 stimulator. In the experimental group, 10 stimuli were delivered with 2-minute inter-stimulus intervals. In the control group, two stimuli were delivered with an 18-minute inter-stimulus interval. The amount of current injected into each cell was set by stepping injections in 0.2nA increments until the cell fired at an average frequency of at least 1Hz. The cell was tested one minute before and one minute after each nerve stimulus. Resting membrane potential was determined by applying a 0.1Hz low pass filter to the traces and taking a 5-second average of the membrane potential 50 seconds before and 50 seconds after a stimulus.

### Statistical analysis

All statistical tests were performed in Prism 10.4.1 (GraphPad). When comparing changes based on flexion phase over repeated stimulation, RM two-way ANOVAs were used to determine significance, followed by multiple comparisons with Tukey’s test. When appropriate, RM one-way ANOVAs followed by Tukey’s test or paired t-tests were used. Alpha value for statistical significance was set at 0.05.

## RESULTS

### Behavioral evidence that sensitization and habituation overlap at some but not all sites in the swim network

The present investigation of how the memories for sensitization and habituation share space in the *Tritonia* swim network began with a behavioral study using a training protocol that produces them in sequence. Previous work had identified three features of *Tritonia*’s swim behavior that can undergo sensitization and/or habituation to repeated swim-inducing tail stimuli: swim onset latency, cycle number, and jump height. However, no previous study had ever tracked all three behavioral components at the same time, as we did here. Animals were divided into experimental and control groups. The experimental group received 10 stimuli of 0.2 mL 4M NaCl to the tail, 2 minutes apart; the control group received the same stimulus 18 minutes apart, the equivalent amount of time as the experimental group (**Fig. 1B**). All stimuli elicited escape swims.

In the experimental group (n = 15), RM one-way ANOVAs found significant effects of training for all three behavioral parameters (**Fig. 1C**; onset latency - F(4.27, 59.78) = 22.32, p <0.0001; jump height - F(4.205, 58.86) = 18.58), p < 0.0001; cycle number - F(5.03, 70.41) = 48.68, p < 0.0001). Multiple comparisons with Tukey’s post-hoc tests from the first to the second stimulus found a statistically significant shortening of onset latency (stim 1: 5.67 ± 1.37s vs. stim 2: 4.10 ± 0.90s, *p* = 0.0005), as well as a statistically significant increase in jump height (stim 1: 99.5 ± 20.9 mm vs. stim 2: 113.0 ± 22.4 mm, *p* = 0.02). In contrast, there was no difference observed in cycle number from stim 1 to stim 2 (stim 1: 5.7 ± 1.0 cycles vs. stim 2: 5.3 ± 1.3 cycles, *p* = 0.98). Thus, sensitization produced by the initial stimulus shortened onset latency and increased jump height, but did not affect cycle number. In the 2-trial control group (n = 17) i.e., no habituation training, paired t-tests found that jump height and onset latency were still sensitized 18 minutes after the initial trial (jump height – stim 1: 87.8 ± 21.6mm vs. stim 2: 101.9 ± 25.5mm, *p* = 0.006; onset latency – stim 1: 6.28 ± 1.80s vs. stim 2: 4.86 ± 0.79s, *p* = 0.0005), and again, cycle number showed no sensitization (paired t-test, stim 1: 5.0 ± 1.6 cycles vs. stim 2: 5.7 ± 1.4 cycles, *p* = 0.16) (**Fig. 1D**). Thus, in the present study, only two parameters of the swim underwent sensitization in response to an initial tail stimulus: onset latency and jump height.

Across the 10 trials of the experimental protocol, the three measured parameters changed in different ways. Onset latency was no longer quickened by stim 4 (stim 1: 5.67 ± 1.37s vs. stim 4: 5.22 ± 0.28s, *p* = 0.76) and remained unchanged from stim 1 by stim 10 (stim 1: 5.67 ± 1.37s vs. stim 10: 6.40 ± 0.46s, *p* = 0.41), displaying a gradual loss of sensitization from repeated stimulation. Jump height was no longer elevated by stim 3 (stim 1: 99.5 ± 20.9mm vs. stim 3: 101.2 ± 24.93mm, *p* > .9999), and showed a significant decrease by stim 7 (stim 1: 99.5 ± 20.9mm vs. stim 7: 75.0 ± 16.9mm, *p* = 0.004), that was even further decreased by stim 10 (stim 1: 99.5 ± 20.9mm vs. stim 10: 65.3 ± 17.7mm, *p* = 0.0006). Thus, jump height’s initial sensitization was gradually replaced by habituation. Cycle number showed a significant decrease by stim 4 (stim 1: 5.7 ± 1.0 cycles vs. stim 4: 3.9 ± 0.8 cycles, *p* = 0.002), that was even more pronounced by stim 10 (stim 1: 5.7 ± 1.0 cycles vs. stim 10: 2.0 ± 0.7 cycles, *p* < 0.0001) (**Fig. 1C)**. Thus, cycle number undergoes gradually accumulating habituation.

These results demonstrate that sensitization and habituation learning did not equally affect all the behavioral parameters of the escape swim, suggesting that the memories for these two forms of learning occupy somewhat independent sites in the network.

### Large-scale optical recording reveals sensitization and habituation act in a phase-specific manner to raise and lower the number of participating dorsal flexion neurons

In a previous study [26], we demonstrated that sensitization learning in *Tritonia* leads to an expansion of the number of reliably bursting neurons within the escape swim network. Neurons that were initially silent or inconsistently active during the first fictive swim became reliable bursters during the second and/or third fictive swim. This expansion of the number of reliably participating neurons represents a unique network component of learning, and we suggested in that prior study that the behavioral correlate of this network expansion was the sensitized jump height. After we demonstrated in the previous section that continued training results in habituation of jump height, here we tested whether the number of reliably participating neurons contracted with repeated stimulation. To do this, we utilized large-scale voltage-sensitive dye (VSD) recordings to capture the neuronal activity of pedal flexion neurons in isolated brains to see if habituation caused neurons to drop out of the motor program with repeated stimulation. The experimental group received 10 fictive swim stimuli delivered to the contralateral pedal nerve 3 (PdN3) 2 minutes apart, while the control group received 2 stimuli delivered 18 minutes apart, mirroring our behavioral protocol. We tracked how many neurons burst during a motor program and if they were dorsal flexion neurons (DFNs) or ventral flexion neurons (VFNs) (**Fig. 2A-B)**. To minimize bleaching of the dye, we did not optically record when stims 5, 7, and 9 were delivered.

**Figure 2.**
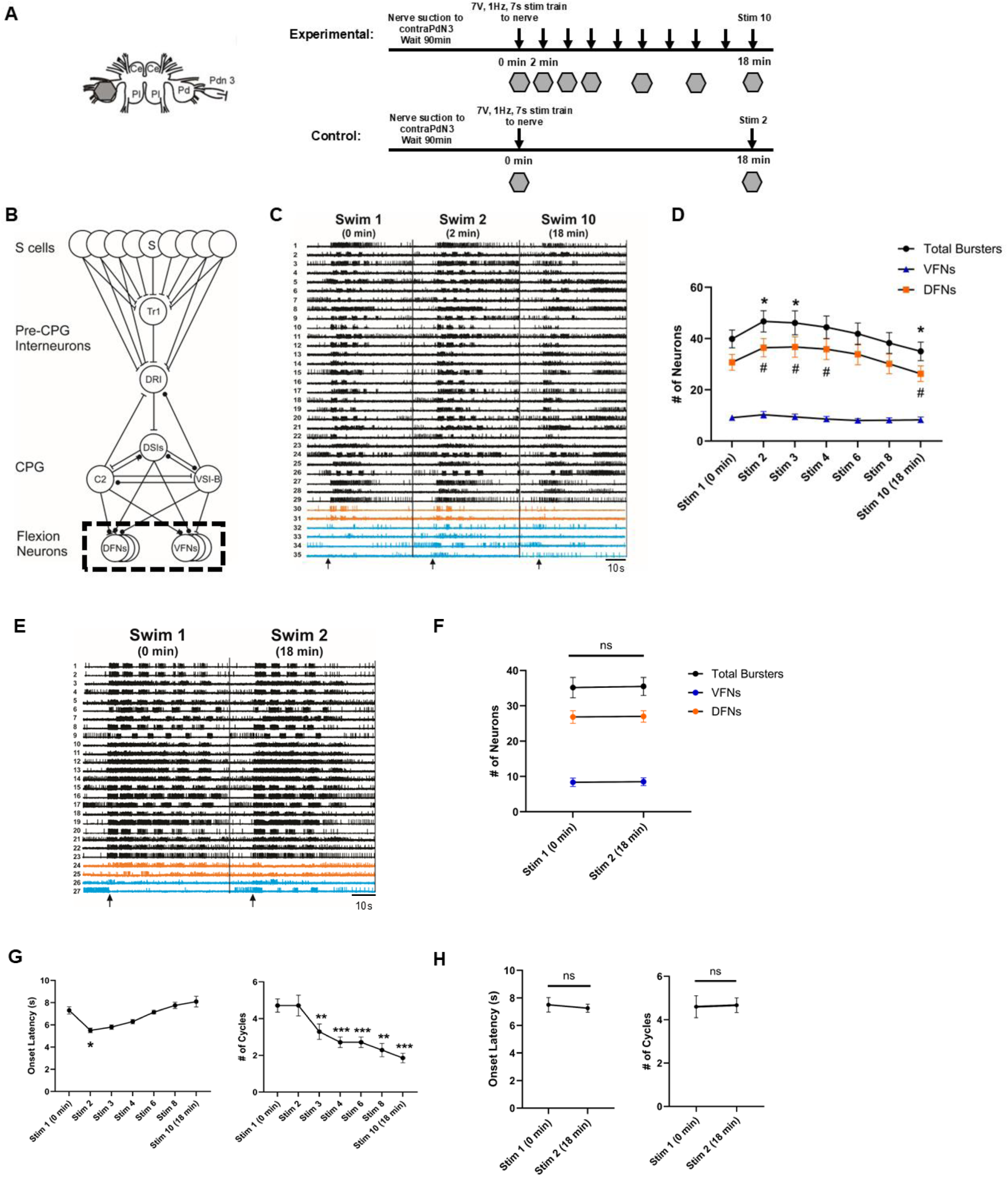
Non-associative learning produces a phase-specific change in neuronal recruitment. **A)** *Left*, illustration of isolated *Tritonia* CNS, showing a grey hexagon over the left dorsal pedal ganglion where activity is captured by the photodiode array, along with a suction electrode attached to the contralateral Pedal Nerve 3. Ce = cerebral, Pl = pleural, Pd = pedal. *Right*, Protocol for isolated CNS voltage-sensitive dye (VSD) recordings. Grey hexagons indicate the times where nerve stimulus was delivered while optically recording with VSDs. **B)** Circuit diagram of *Tritonia* SMP – black dashed box indicates the portion of network captured via VSD imaging - the phasic flexion neurons. **C)** Representative example of how neuronal participation changes with repeated stimulation (black arrow indicates when the stimulus was delivered). Recordings from all 7 optical recordings were concatenated before spike sorting. Black traces are neurons that burst in every fictive motor program. Orange traces are neurons that started in the network but ultimately left the network by the last recording. Blue traces are neurons that were not bursting at stim 1, that joined on stim 2, and left by stim 10. Breaks between recordings shown with black line. **D)** Change in neuronal participation with repeated stimulation (n = 7 preparations). The total number of bursters increases under sensitization and decreases under habituation, however, all of the change is in the number of dorsal neurons. Significance for total bursters compared to stim 1 indicated with * (*p* < 0.05). Significance for DFNs compared to stim 1 indicated with # (*p* < 0.05). **E)** Representative example of how neuronal participation does not change in the control condition. Same color scheme as **C.** Stimulus indicated by black arrow. Black line in the center designates break between recordings. **F)** No change in neuronal participation on either phase in control condition (n = 6 preparations). **G)** Changes in onset latency and cycle number in the experimental preparations described demonstrates the development of sensitization by stim 1 and habituation by stim 10. Significant differences compared to stim 1 shown with * (*p* < 0.05), ** (*p* < 0.01), or *** (*p* < 0.001). **H)** Lack of changes in onset latency and cycle number in the control preparations shows sensitization did not persist in isolated brains at 18 minutes. Error bars represent SEM.

We confirmed that the experimental preparations (n = 7) underwent non-associative learning by tracking onset latency and cycle number in each recording and identified significant differences in both onset latency (F(1.779, 10.68) = 15.23, *p* = 0.0009) and cycle number (F(2.774, 16.64) = 25.00, *p* < 0.0001) via RM one-way ANOVAs (**Fig. 2G)**. Tukey’s post-hoc tests identified onset latency shortened from stim 1 to stim 2 (stim 1: 7.30 ± 0.85s vs. stim 2: 5.51 ± 0.53s, *p* = 0.02), and cycle number habituated from stim 1 to stim 10 (stim 1: 4.7 ± 0.9 cycles vs. stim 10: 1.9 ± 0.7 cycles, *p* = 0.0003), consistent with the development of an initial sensitization by stim 2 and eventual habituation by stim 10 in these preparations. In our control preparations (n = 6), paired t-tests confirmed that both onset latency (stim 1: 7.50 ± 1.29s vs. stim 2: 7.26 ± 0.72s, *p* = 0.62) and cycle number (stim 1: 4.5 ± 1.0 cycles vs. stim 2: 4.7 ± 0.8 cycles, *p* = 0.61) were unchanged (**Fig. 2H)**, verifying that these preparations were no longer sensitized after 18 minutes.

In the experimental group, we observed a significant change with repeated stimulation in the number of bursters across phases (RM two-way ANOVA, F(12, 108) = 12.00), *p* < 0.0001) (**Fig. 2C-D)**. Tukey’s post-hoc tests afterwards revealed the total number of bursters increased from stim 1 to stim 2 (stim 1: 39.9 ± 9.2 neurons vs. stim 2: 46.7 ± 11.0 neurons, *p* = 0.005). This increased overall burster network size remained until stim 6 (stim 1: 39.9 ± 9.2 neurons vs. stim 6: 41.9 ± 11.2 neurons, *p* = 0.51). By stim 10, we observed a slight decrease in the overall network size, (stim 1: 39.9 ± 9.2 neurons vs. stim 10: 35.0 ± 9.6 neurons, *p* = 0.02). Notably, however, this change was specific to the dorsal phase. From stim 1 to 2, the number of DFNs significantly increased (stim 1: 30.7 ± 8.2 neurons vs. stim 2: 36.4 ± 9.4 neurons, *p* = 0.005). The number of DFNs remained increased until stim 6 (33.9 ± 10.8 neurons, *p =* 0.19), and was significantly decreased by stim 10 (26.3 ± 8.0 neurons, *p* = 0.02). In comparison, there was no significant change in the number of VFNs from stim 1 to stim 2 (stim 1: 9.1 ± 2.8 neurons vs. stim 2: 10.3 ± 3.5 neurons, *p* = 0.12) or from stim 1 to stim 10 (stim 1: 9.1 ± 2.8 neurons vs. stim 10: 8.3 ± 2.9 neurons, *p* = 0.13). Thus, the alterations in the size of the bursting network were solely due to the recruitment of and loss of DFNs.

By comparison, in our control group, which as mentioned did not undergo habituation training, we observed no significant change in the total number of neurons participating in the motor program (RM two-way ANOVA, F(1,5) = 0.357, *p* = 0.58), (stim 1: 35.2 ± 7.0 neurons vs. stim 2: 35.5 ± 6.5 neurons, *p* = 0.89) (**Fig. 2E-F)**. This demonstrated that the recruitment of neurons after a sensitizing stimulus had faded within 18 minutes in isolated brains, and that the loss in neurons observed in the experimental group was specifically the result of habituation learning.

These findings demonstrate that both sensitization and habituation alter the size of the efferent network. However, they both do so in a phase-specific manner, recruiting and dropping out neurons on the dorsal phase. This makes network size alterations an insufficient explanation for changes in jump height, as the jump is powered by the first ventral flexion.

### Sensitization and habituation do not equally alter DFN and VFN burst intensity

The alternating bursts of activity in the VFNs and DFNs of the pedal ganglion generate the ventral and dorsal body flexions of the swim [31]. Thus, changes in the intensity of their firing during a motor program may explain the changes in the behavior. The jump occurs during the first ventral flexion, so we took advantage of our large-scale VSD recordings described in the previous section to assess how sensitization and habituation learning changed the burst intensity on the first flexion phase. We restricted our analysis to flexion neurons that burst on the first flexion of every fictive motor program.

In the experimental group, we found that the first cycle burst intensity in both DFNs and VFNs changed significantly with repeated stimulation (RM two-way ANOVA, F(6, 72) = 2.636, *p* = 0.02) (**Fig. 3A-B)**. Tukey’s post-hoc tests showed that the VFN burst intensity increased significantly from stim 1 to stim 2 (stim 1: 31.2 ± 8.3 spikes vs. stim 2: 37.9 ± 9.9 spikes, *p* = 0.001), consistent with the sensitization-driven enhancement in jump height seen behaviorally. In contrast, DFN burst intensity remained unchanged (stim 1: 20.4 ± 4.1 spikes vs. stim 2: 19.2 ± 6.6 spikes, *p* = 0.90), indicating that sensitization selectively strengthens ventral phase output. VFN burst intensity decreased significantly by stim 8 (stim 8: 20.7 ± 3.9 spikes, *p* = 0.02), and was further decreased by stim 10 (19.6 ± 7.8 spikes, *p* = 0.005), mirroring behavioral habituation of jump height. Notably, repeated stimulation also significantly reduced DFN burst intensity by stim 8 (10.8 ± 2.5 spikes, *p* = 0.02), that was also further reduced by stim 10 (9.0 ± 1.6 spikes, *p* = 0.005), demonstrating that habituation weakens the first cycle activity in both phases.

**Figure 3.**
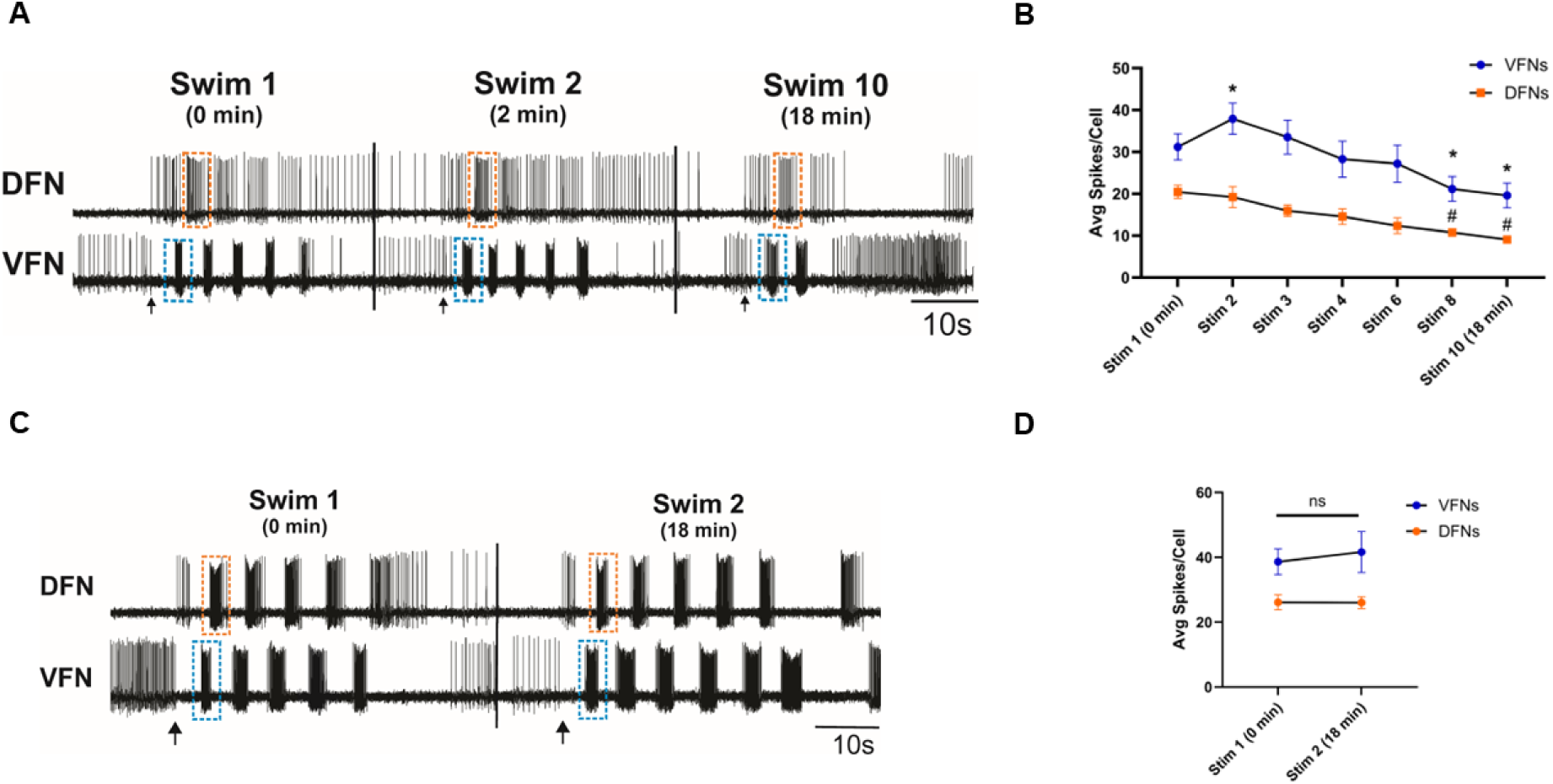
Sensitization and habituation learning differentially modify first flexion burst intensity. **A)** Representative optical recordings using VSDs of a reliably bursting DFN & VFN that participated in every fictive swim program in the experimental preparations. Orange and blue dotted boxes are placed to highlight the first dorsal and ventral flexion, respectively, demonstrating the increase in VFN firing on stim 2 and the decrease in VFN & DFN firing on stim 10. Stimulus marked by black arrow. Black lines indicate breaks between recordings. **B)** Changes in average burst intensity per recording in the experimental group. Significant differences in VFN firing compared to stim 1 indicated with * (*p* < 0.05). Significant differences in DFN firing compared to stim 1 indicated with # (*p <* 0.05). **C)** Representative optical recordings using VSDs of a DFN and VFN that participated in both fictive swim programs in the control preparations. Same coloring scheme as **A**. Stimulus indicated by black arrow. Black line indicates the end of the first recording. **D)** No change in the first flexion burst intensity of VFNs or DFNs in the control preparations. Error bars represent SEM.

In our control experiments (two stimuli, 18 minutes apart), we found no significant change in first flexion burst intensity in either DFNs or VFNs (RM two-way ANOVA, F(1,10) = 0.8238, *p* = 0.39) (**Fig. 3C-D)**, which demonstrated that the changes in burst intensity on both phases identified in the experimental group were the result of the learning.

These findings reveal distinct sites of plasticity for sensitization and habituation. Sensitization enhances VFN burst intensity, directly correlating with the behavioral increase in jump height, without affecting DFN burst intensity. In contrast, habituation decreases burst intensity in both phases, erasing the sensitization-driven enhancement in VFN burst intensity, and producing a decrement in DFN burst intensity. Thus, while both forms of learning can affect the same network feature (burst intensity), they do not act on each flexion phase in the same manner. This also helps explain how jump height can change without any change in the number of participating VFNs – the burst intensity of the VFNs is what changes.

### Sharp electrode recordings reveal that non-associative learning differentially alters excitability depending on flexion phase

The observed different changes in first flexion burst intensity under sensitization and habituation learning led us to investigate what potential cellular or intrinsic properties may underlie these changes. We recorded DFNs and VFNs in the pedal ganglion of isolated *Tritonia* CNS with intracellular electrodes and tracked changes in their intrinsic excitability (as determined by 5-second constant current injections) before and after fictive swims, as well as their first cycle burst intensity. We hypothesized that the excitability of these neurons would change in similar ways with their burst intensity changes during the motor program: VFNs would increase excitability under sensitization and decrease under habituation, and DFNs would only decrease under habituation. As in our VSD experiments, our experimental group had 10 stimuli delivered to the contralateral PdN3 2 minutes apart, and our control group had 2 stimuli delivered 18 minutes apart.

To confirm the presence of sensitization and habituation in the experimental group, and the lack of either form of learning in the control group, we also tracked onset latency and cycle number in these recordings (**Fig. 4D**). In the experimental group, RM one-way ANOVAs identified significant changes in onset latency (F(2.921, 23.37) = 7.535, *p* = 0.001) and cycle number (F(2.528, 20.22) = 14.81, *p* < 0.0001). Tukey’s post-hoc tests identified a decrease in onset latency at stim 2 (stim 1: 7.83 ± 1.65s vs. stim 2: 5.95 ± 1.22s, *p* = 0.04), and no change in cycle number (stim 1: 3.2 ± 0.9 cycles vs. stim 2: 3.1 ± 0.8 cycles, *p* = 0.99), confirming the development of sensitization in these intracellular recordings and consistent with our behavioral experiment. At stim 10, we observed a decrease in cycle number (stim 10: 1.3 +/− 0.5 cycles, *p* = 0.03) and a return to naïve latency (stim 10: 8.33 ± 2.10s, *p* = 0.99), confirming the development of habituation in these recordings. In our control preparations (**Fig. 4E)**, paired t-tests showed onset latency (stim 1: 7.93 ± 1.19s vs. stim 2: 8.14 ± 1.32s, *p* = 0.47) and cycle number (stim 1: 3.8 ± 0.9 cycles vs. stim 2: 3.9 ± 0.9 cycles, *p* = 0.78) displayed no significant changes, which demonstrated that the sensitization learning had faded within 18 minutes, as in our VSD experiments described earlier.

**Figure 4.**
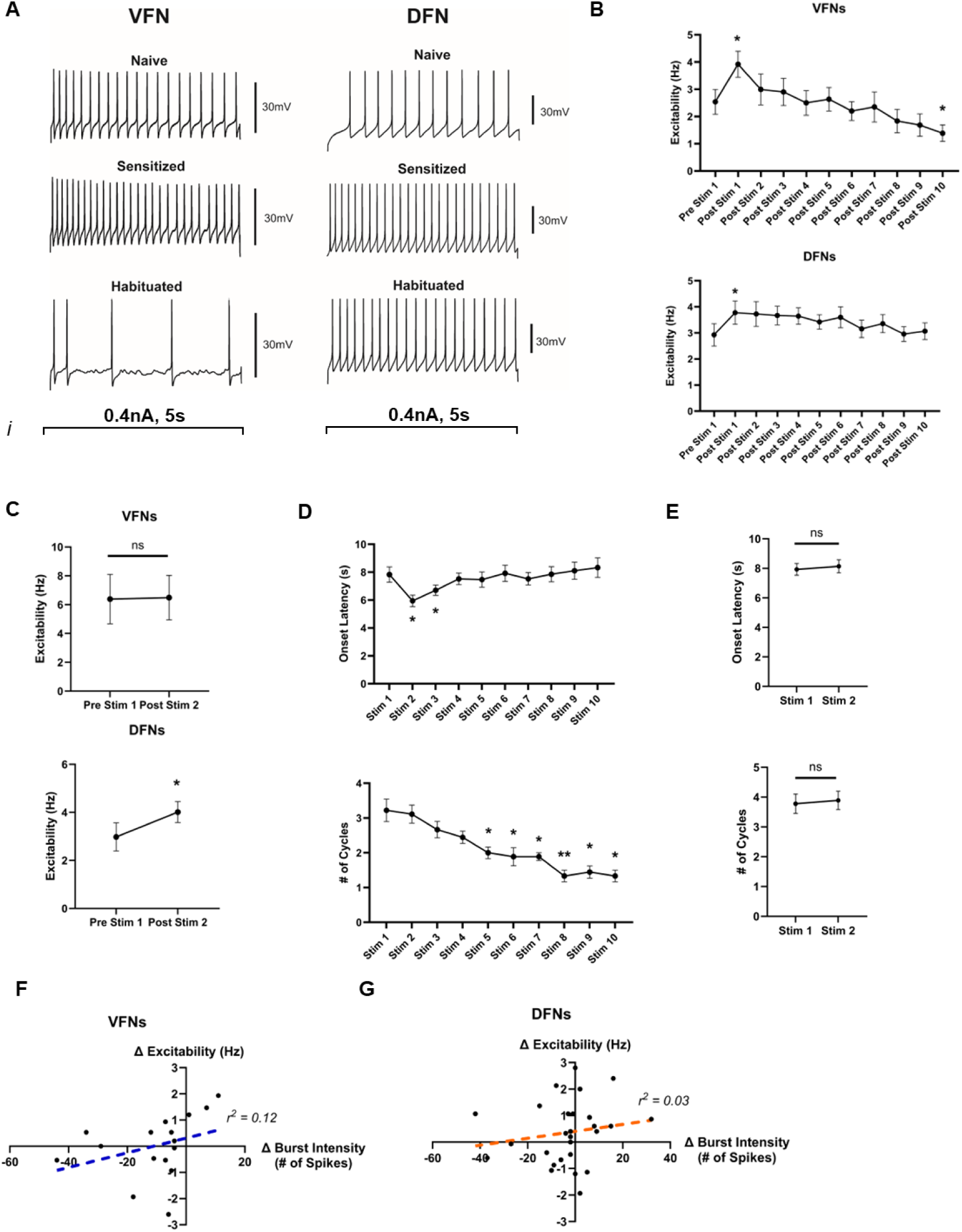
Learning induced changes in excitability are dependent on flexion phase. **A)** Representative examples of intrinsic excitability tests in the same VFN and DFN under naïve, sensitized, and habituated conditions in the experimental group. VFN and DFN excitability increase under sensitization, and VFN excitability decreases under habituation, while DFN excitability does not. **B)** Changes in excitability in all VFNs and DFNs recorded in the experimental groups. * indicates significant difference compared to pre-stim 1 (*p* < 0.05). N = 9 VFNs, 16 DFNs. **C**) Loss of sensitization increase in VFN excitability within 18 minutes in control condition, however DFN excitability remains elevated (* indicates *p* < 0.05). **D)** Validation of development of sensitization and habituation in the experimental group via changes in onset latency and cycle number. Significant differences relative to stim 1 indicated by * (*p* < 0.05), or ** (*p* < 0.01). **E)** Validation of sensitization dissipation in control group via lack of changes in onset latency or cycle number. **F)** Linear regression comparing changes in VFN excitability and burst intensity was not significant (regression line shown with blue dashed line), indicating that increased burst intensity is not caused by increased excitability. **G)** Same finding as **F,** but in DFNs (regression line shown with orange dashed line). Error bars represent SEM.

When looking at potential changes in excitability in the experimental group, repeated measures one-way ANOVAs for the ventral phase excitability (n = 6, F(2.97, 14.85) = 7.755, *p* = 0.002) and the dorsal phase excitability (n = 9, F(4.097, 32.78) = 3.159, *p* = 0.03) were both significant. Tukey’s post-hoc tests showed that after an initial sensitizing stimulus, intrinsic excitability increased for both VFNs (pre-stim 1: 2.54 ± 1.12Hz vs. post-stim 1: 3.92 ± 1.17Hz, *p* = 0.02) and DFNs (pre-stim 1: 2.93 ± 1.28Hz vs. post-stim 1: 3.78 ± 1.34Hz, *p* = 0.01) (**Fig. 4A-B**). After stimulus 10, intrinsic excitability only decreased for VFNs (post-stim 10: 1.39 ± 0.74Hz, *p* = 0.04) – DFNs showed no significant change in excitability from naïve (post-stim 10: 3.03 ± 0.35Hz, *p* = 0.99) (**Fig. 3B**). In our control group, with stimuli 18 minutes apart, paired t-tests revealed no change in excitability in VFNs (n = 6, pre-stim: 6.38 ± 1.71Hz vs. post-stim 2: 6.49 ± 1.54Hz). Interestingly, however, we found that excitability remained increased in DFNs in these control preparations (n = 9, paired t-test, pre-stim: 2.98 ± 1.75Hz vs. post stim 2: 4.01 ± 1.3Hz, *p* = 0.04) (**Fig. 4C**). VFN excitability therefore tracked with our hypothesis of an increase with sensitization and a decrease with habituation, as well as remaining unchanged in preparations that did not display sensitization. DFN excitability, however, did not follow our hypothesis, increasing with sensitization and only returning to naïve as the preparation became habituated. Even in control preparations that displayed a loss of sensitization, DFN excitability was still increased, suggesting this form of sensitization learning-induced change persists beyond the loss of fictive sensitization.

We then investigated if changes in excitability in these flexion neurons were correlated with changes in their first cycle burst intensity, to determine if the changes to intrinsic excitability can explain the changes to motor program activity previously described. However, linear regression models comparing excitability changes with first cycle burst intensity changes on both ventral (**Fig. 4F**, r^2^ = 0.12, F(1,14) = 1.852, *p* = 0.19) and dorsal phase (**Fig. 4G**, r^2^ = 0.03, F(1, 25) = 0.727, *p* = 0.40) were both not significant. This demonstrated that changes in intrinsic excitability cannot directly explain the altered motor program activity on either phase.

These results indicate that excitability changed in both ventral and dorsal flexion neurons when non-associative learning occurred, but these changes are not consistent across flexion phase and do not explain changes to the motor program. VFN and DFN excitability increased under sensitization, but only VFN excitability decreased under habituation. DFN excitability also remained elevated even when the preparation no longer demonstrated a sensitized fictive behavior, potentially representing a persistent component of the sensitization learning that outlasts the behavioral change. However, changes in excitability on either phase were not correlated with changes in burst intensity, which indicates that learning induced changes in burst intensity are likely driven by changes in synaptic input, not intrinsic excitability.

### Ventral and dorsal phases use distinct mechanisms to alter intrinsic excitability

While the changes in the first cycle burst intensity could not be explained by the changes in excitability, we were interested in how excitability did not change consistently between the dorsal and ventral phases, as it suggested different mechanisms for altering intrinsic excitability during learning. We focused on two previously studied cellular mechanisms for excitability – changes in resting membrane potential (RMP) and changes in spike frequency adaptation (SFA). Changes in RMP have been demonstrated in neurons with altered excitability in multiple different species, suggesting it is a common conserved cellular mechanism [32, 33]. SFA is a membrane property observed in many neurons where sustained firing leads to a progressive reduction in instantaneous spike frequency, often mediated by Ca^2+^-activated K^+^ channels [34]. A reduction in SFA would increase excitability, and an increase in SFA would decrease excitability. To test for changes in these properties, we re-analyzed the data from the intracellular experiments described earlier.

RMP for VFNs in the experimental group was unchanged with repeated stimulation relative to the pre-stim resting potential (RM one-way ANOVA, F(2.093, 8.372) = 0.718, *p* = 0.52). However, the RMP of DFNs in the experimental group did change (RM one-way ANOVA (F(2.751, 22.01) = 7.456, *p* = 0.002), and Dunnet’s post-hoc tests revealed DFN RMP was depolarized after the first stimulus (post-stim 1: +8.9 ± 7.0 mV, *p* = 0.03), and remained depolarized with repeated stimuli (post-stim 10: +9.8 ± 6.7 mV, *p* = 0.04) (**Fig. 5A-B**). In our control group, paired t-tests revealed no significant change in RMP in VFNs relative to the pre-stim RMP (post-stim 2: +3.3 ± 4.4 mV, *p* = 0.12), and interestingly, a depolarized RMP in the DFNs (post-stim 2: +3.6 ± 2.8 mV, *p* = 0.005) (**Fig. 5C**). A simple linear regression between changes in intrinsic excitability and RMP in VFNs (r^2^ = 0.01, F(1,14) = 0.1525, *p* = 0.70) was not significant, but it was significant between excitability and RMP in DFNs (r^2^ = 0.23, F(1,24) = 7.089, *p* = 0.01) (**Fig. 5D)**. Thus, RMP does not appear to change in VFNs during non-associative learning, ruling it out as a mechanism for driving changes in VFN excitability. RMP was depolarized in DFNs, however, under sensitization, and remained so under habituation. This suggests that RMP does contribute to the increased excitability of the DFNs, although it does not appear to be the only mechanism driving changes in DFN excitability. This sensitization-led depolarization in DFN RMP also persists even after the sensitization learning fades, suggesting an additional role for DFN RMP depolarization.

**Figure 5.**
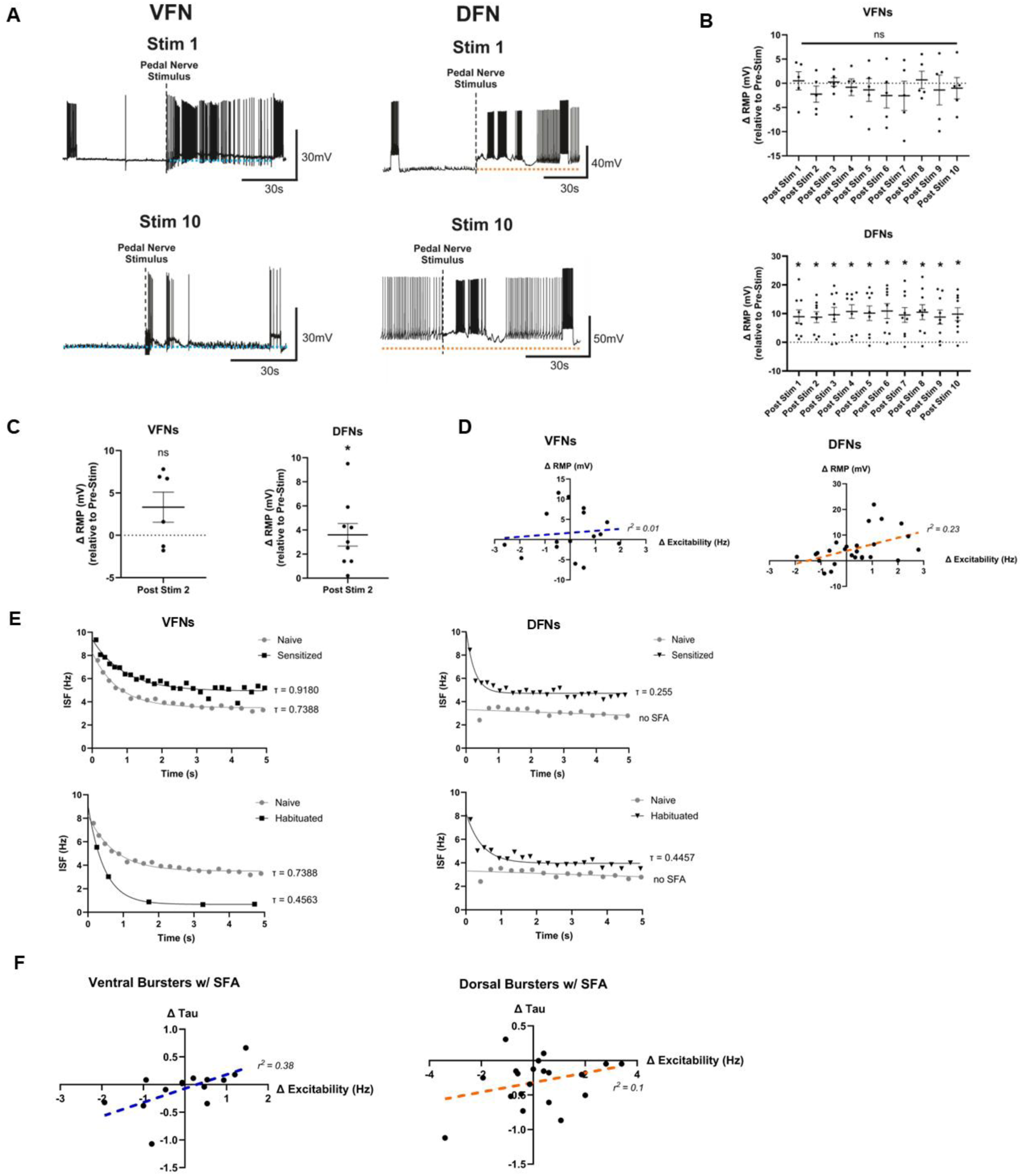
Intrinsic excitability is changed via different mechanisms depending on flexion phase. **A)** *Left*, Representative example of VFN in the experimental group that displayed increased excitability to the 5s test pulse after the first nerve stimulus (Top), and decreased excitability by stim 10 (Bottom), as shown in the group data in **B**. These changes were not accompanied by any change in RMP (as indicated by the blue dashed line (Top). *Right*, Representative example of a DFN in the experimental group that displayed increased excitability to the 5s test pulse after the first nerve stimulus (Top), which was not decreased from naïve by stim 10 (Bottom), as shown in the group data in **B**. These changes in excitability were accompanied by a depolarized RMP after the first nerve stimulus (Top), as indicated by the orange dashed line, which was still depolarized by stim 10 (Bottom). **B)** Group data for the effects shown in **A**. Significant differences compared to pre-stim 1 indicated with * (p < 0.05). **C)** *Left,* No change in VFN RMP in the control condition. *Right*, DFN RMP remained depolarized during the follow-up stimulus in the control condition. **D)** *Left*, Non-significant linear regression (regression line is blue dashed line) indicates no relationship between changes in excitability and changes in RMP in VFNs. *Right*, Significant linear regression (regression line is orange dashed line) indicates relationship between changes in excitability and changes in RMP in DFNs. **E)** Graphs of instantaneous spike frequency (ISF) of the VFN and DFN excitability tests displayed in Fig. 4A (experimental group). One phase decay curve fit to each graph and Tau value calculated and displayed. *Left*, VFN neuron displays reduced SFA (higher Tau value) under sensitization, and increased SFA (lower Tau value) under habituation. *Right*, Increases and decreases in excitability in DFN have no relationship to Tau values. **F)** *Left*, Significant linear regression (regression line is blue dashed line) indicating a relationship between changes in excitability and changes in SFA in VFNs. *Right*, Non-significant linear regression (regression line is orange dashed line) indicating no relationship between changes in excitability and changes in SFA in DFNs. Error bars represent SEM.

**Figure 6.**
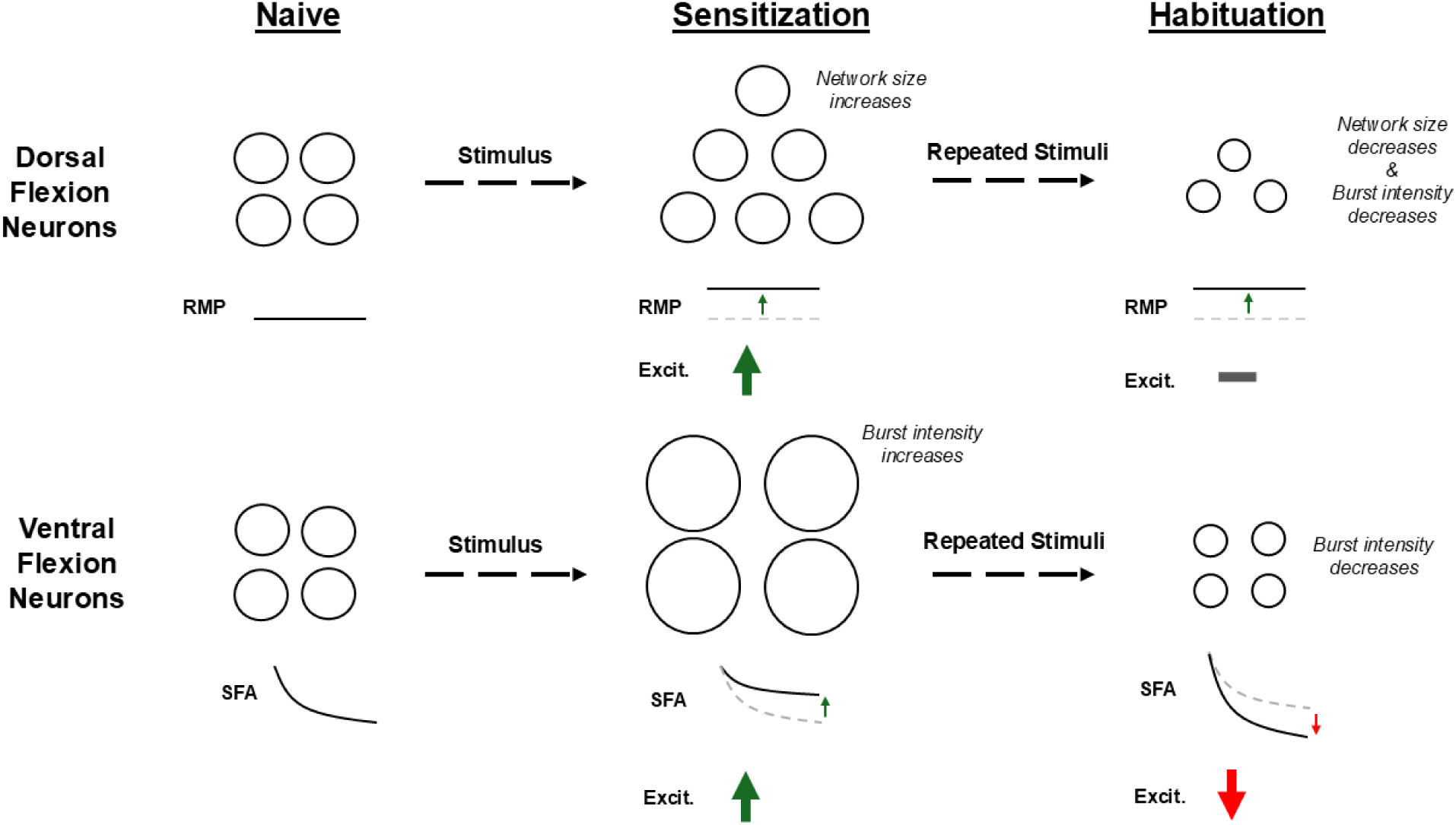
Summary of findings. Sensitization and habituation both make different alterations to *Tritonia*’s escape network, modifying distinct cellular properties and differentially altering activity on different phases. With sensitization, the number of DFNs increases (indicated by the increased number of circles), while the intensity of VFN firing increases (indicated by the increased size of the circles). Different cellular mechanisms are involved for each phase. Under habituation, the number of DFNs decreases and they fire less intensely (indicated by the reduced number of circles and the smaller size of the circles), while the VFNs again do not change participation number and only fire less intensely. DFNs and VFNs use different cellular mechanisms to achieve these changes – DFNs raise RMP under sensitization, and keep it elevated under habituation, while VFNs reduce SFA under sensitization and increase it under habituation.

We also sought to determine if changes in SFA drove changes in excitability in either phase. To do this, we determined the instantaneous spike frequency (ISF) of both VFNs and DFNs during the excitability tests done in the previous section and determined if they exhibited SFA by using the metric noted in [35]. A majority of DFNs and VFNs at rest exhibited SFA. In these SFA-displaying neurons, we quantified the rate of one-phase exponential decay (Tau) (**Fig. 5E)**. In VFNs, changes in excitability were positively correlated with changes in Tau—a slower rate of ISF decay (higher Tau) corresponded to increased excitability (simple linear regression, F(1,12) = 7.308, r^2^ = 0.38, *p* = 0.02) (**Fig. 5F**). Functionally, this implies that a given synaptic input would evoke more spikes due to reduced SFA. In contrast, DFNs showed no significant correlation between changes in excitability and Tau (simple linear regression, F(1,18) = 2.006, r^2^ = 0.10, *p* = 0.17) (**Fig. 5E-F**). Therefore, DFNs do not change excitability by modifying SFA.

These results suggest that VFNs regulate excitability primarily through modifications to SFA, while DFNs rely on changes in RMP. Thus, the two neuron types employ distinct mechanisms to modulate their intrinsic excitability under sensitization and habituation.

## DISCUSSION

While there is growing interest in the multiple mechanisms involved in balancing overlapping memory formation [36, 37], there are limited avenues for precisely interrogating how different modifications are utilized to store information to help to keep different memories apart within the same small network space. Even in simple marine invertebrates, a single memory can produce multiple distributed alterations within a network [38]. Here, we were successfully able to utilize *Tritonia*’s escape swim network to reveal how two competing forms of non-associative learning (sensitization and habituation) use distinct network alterations to differentially modify multiple components of the animal’s escape behavior. Even though these two memories are encoded within the same network and can interfere with each other, they both utilize a combination of shared and distinct sites of plasticity. Each memory is therefore utilizing a strategy of multiple network alterations, some shared and some unique, to change behavior – a potential method for allowing multiple memories to co-exist within the same network.

### Sensitization and habituation modify escape behavior in different ways

Many previous studies have explored behavioral non-associative learning in *Tritonia’s* escape swim [22, 24, 25, 39], but this is the first study to track three behavioral components (onset latency, cycle number, and jump height) undergoing sensitization and habituation in the same animals. These three components did all change, but not in the same manner, showing that distinct components of the same behavior are being differentially modified by the same repeated trial protocol. Sensitization and habituation therefore appear to modify multiple distinct components of the same swim network, as has been speculated in prior work [39, 40]. In this study, we identified jump height as another feature of the escape swim that can undergo habituation and observed only a return to naïve onset latency with repeated stimulation. These results suggest that habituation learning destructively interferes with sensitization in this behavioral parameter. Onset latency, however, is a parameter that initially sensitizes and loses this sensitization during habituation training. The finding that the control procedure, two single swim trials separated by 18 minutes, produces a persisting onset latency sensitization presents two possibilities. Latency sensitization and habituation could be encoded via different sites of plasticity, which results in destructive interference in the behavioral output but both memories’ latency plasticity persists within the neural network. Alternatively, latency plasticity is shared between sensitization and habituation, and thus these behavioral results demonstrate destructive interference between the two forms of learning.

Prior work has demonstrated that onset latency can remain sensitized even when cycle number is habituated, supporting the argument that the two memories do not use all the same sites across behavioral parameters [24, 40]. While we only observed cycle number habituation in our experiments, it has also been shown that cycle number can sensitize with a weaker, nonmaximal training stimulus [39]. Some of these differences may be the result of different batches of animals collected in different years, as well as from different collection points. However, since jump height appeared to be a definitive site of destructive interference between the two forms of learning in our behavioral experiments, we then sought to identify a neural correlate for this behavior that could be investigated as a site of interfering plasticity between two memories.

### Network size modification during learning was specific to one phase

Two ways to modify network function in learning are to either change the vigor with which existing members of the active network fire, or to change the number of participants [2]. Changes in numbers of participants has previously been tracked with large scale recording methods [41, 42], which have also been applied to study learning in *Tritonia* and *Aplysia* [17, 43]. During the *Tritonia* swim motor program, many neurons participate variably, firing on some cycles of the swim and not others [30]. With sensitization learning, many variable and non-bursters become reliable bursters, increasing the overall size of the swim network, presumably to increase overall swim strength [26]. Serotonin release from the DSIs, the serotonergic neurons within the CPG, has been shown to drive this network expansion [26]. We were interested in determining if this network expansion in sensitization would be followed by network contraction in habituation, as a possible explanation for jump height habituation. It is difficult to assess variable or reliable participants in a habituated preparation, as the network is often only producing one or two swim cycles in total, hence our reliance on total network size to focus on if bursters were shifting to becoming non-bursters with repeated stimulation.

In this study, we identified network expansion under sensitization that gave way to network contraction under habituation. This represents a network plasticity site where the two memories interfere. However, when we broke down the changes by flexion phase, we identified that all the changes came from recruitment of or loss of DFNs. There was no significant change in the share of VFNs with repeated stimulation. This suggested that a different element of the network is specifically responsible for the changes in jump height. It is possible that the changes in the number of DFNs would alter the strength of the dorsal flexion, but we currently lack the capacity to measure this change in the intact animal.

### Sensitization and habituation are not the inverse of one another at the cellular level

A potential neural correlate of *Tritonia*’s jump height was the burst intensity on the first ventral phase, which we hypothesized would increase under sensitization and decrease under habituation. This is precisely what we observed with our VSD experiments. Since the ventral and dorsal phases are very similar in duration and visually appear as the inverse of one another, we hypothesized that the first dorsal phase would also display increased burst intensity under sensitization and decreased intensity under habituation. However, the finding that the first dorsal phase burst intensity did not increase under sensitization, but did decrease under habituation, demonstrates that sensitization and habituation are utilizing different mechanisms and modulation to change behavior. Each memory alters burst intensity, but sensitization does so in a phase-specific way, whereas habituation does so in a phase-general way. This result also helps explain how jump height can sensitize without new VFNs joining the network - the burst intensity of the VFNs is increased. Meanwhile, the DFNs may not need to fire more intensely under sensitization to strengthen the dorsal flexion, because more of them are recruited into the network. The cumulative result of these changes would be a strengthened swim in both flexion phases, but via different mechanisms. With repeated stimulation, cycle number quickly starts habituating, but the flexion neurons stay involved and continue firing vigorously, only decreasing their burst intensity after many repeated stimuli. This suggests two possibilities – either habituation learning is slowly erasing sensitization in these neurons, or sensitization and habituation learning are coexisting in these neurons, effectively cancelling each other out in the behavior until eventually sensitization’s plasticity fades or is overwhelmed by habituation’s plasticity. This would produce behavioral interference via independent sites of plasticity.

Other work has shown that sensitization and habituation are distinct forms of learning that can often co-exist and overlap within different parameters of the same animal’s behavior [44, 45]. In *Tritonia*, the bulk of both behavioral and cellular investigations into learning have focused on the mechanisms of sensitization, but this study shows that habituation in *Tritonia* involves unique pathways and mechanisms that have been previously unexplored. Future work to identify what cellular and molecular mechanisms drive habituation of other parameters of *Tritonia*’s swim, like cycle number, should therefore also anticipate distributed sites of plasticity within the network, including sites that have not been previously characterized.

### Different sites of plasticity may store different information about behavioral changes

Since the ventral and dorsal flexion neurons are efferent neurons receiving input from the CPG, there are two logical mechanisms for any increase or decrease in their activity – either the input they receive has changed, and/or their intrinsic properties have changed. The historical literature on learning-induced changes across various species has largely focused on changes to synaptic strength [46–48]. However, the effects of learning on a neuron’s intrinsic membrane properties have also been explored for decades, and have often been identified in tandem with changes to synapses [38, 49, 50]. There is a growing appreciation in the literature of the importance of intrinsic excitability and other membrane properties in recruiting neurons into neural networks [1, 2, 51]. Since *Tritonia*’s SMP CPG contacts a large population of efferent neurons to produce the behavior, it is plausible that efferent output could be altered by changing the excitability of the efferent neurons.

Our intracellular recordings revealed a sensitization-induced increase in intrinsic excitability on both dorsal and ventral phases, but a habituation-induced decrease on only the ventral phase. The changes in DFN excitability appear to be driven at least in part by a depolarization in RMP. This depolarization of RMP in DFNs persists even after onset latency sensitization has faded with time or been erased as habituation develops in other behavioral parameters. This is an example of how sensitization learning plasticity can co-exist with habituation learning plasticity. We do not currently know what drives the depolarized RMP, but we hypothesize it could be due to CPG neuromodulation or rapid temporally summating EPSPs. Future work is needed to test these possibilities. The VFNs do not undergo any change in RMP with learning – the changes in their excitability appear to be primarily driven by changes in the intensity of spike frequency adaptation. Modifying SFA to mediate excitability is a common mechanism across species [52] and is even used within the *Tritonia* escape swim – C2, a peptidergic neuron within the CPG, becomes more excitable during the fictive escape swim via a loss of SFA due to serotonin release from the DSIs [53]. We do not currently know what drives the changes in SFA in VFNs. We hypothesize that neuromodulation from the CPG is responsible for this modulation. Future work is needed to test this hypothesis. Thus, different memories can act on the same network via the same site of plasticity, efferent neuron excitability, through different cellular mechanisms.

We did not see a correlation between a change in burst intensity and a change in intrinsic excitability on either flexion phase. This suggests that the likely mechanism directly driving changes in burst intensity during a flexion is upstream, coming from summation of synaptic inputs from the respective phase CPG neurons. The primary ventral CPG neuron, VSI-B, is known to increase its output onto VFNs immediately after a fictive swim [54], which could drive the increase in VFN burst intensity under sensitization. We hypothesize that VSI-B is decreasing its output onto the VFNs with successive stimuli, and that the DSIs and C2 are doing the same to DFNs as well. Future work will be needed to test this hypothesis.

Many recent studies have identified increased excitability as a crucial mechanism involved in recruiting neurons into a memory trace and effecting the corresponding behavior [51, 55]. While we observe learning-induced changes in the animal’s behavior, changes in the intrinsic excitability of the efferent neurons producing this behavior, and increased activity in these neurons during the fictive behavior, the specifics of our preparation allow us to see that the latter two are not directly correlated. How excitable a flexion neuron is at rest does not correlate with how vigorously it participates in the motor program. So, what do the changes in excitability represent? The changes in excitability may instead be involved with a different component of the behavior. Habituation’s phase-specific effects on excitability may also be relevant, as there are more DFNs than VFNs in the network [28], and our optical recordings demonstrated that VFNs on average fire more vigorously compared to DFNs. DFNs may be recruited under sensitization via increased excitability and may remain as weak participants under habituation via elevated RMP and a lack of reduced excitability. This may serve as network protection against a critical loss of DFN participation that could prevent the production of a dorsal flexion and compromise the escape swim.

These results demonstrate that each site of plasticity may store different information or be responsible for a different dimension of the overall behavior. This insight is only possible because of the simplicity of our model and the direct relationship between neural activity and quantifiable motor behavior.

### Sequential memories in the same network are using different strategies to encode distributed sites of plasticity

We have shown here that sensitization and habituation memories in *Tritonia*’s escape swim network are employing different strategies to produce distributed sites of plasticity throughout the escape network. As the escape swim is a stereotyped behavior with several behavioral parameters, driven by a small network, it follows that a possible method to modify one feature of the behavior and not another is to make distinct modifications to specific features of the network. Our behavioral results showed sensitization and habituation acting differently across the various parameters, but we did not observe behavioral overlap of these two memories as we have in previous studies [24, 40].

Even when these two memories produce behavioral interference, however, we have shown demonstrable evidence of distributed organization of these two memories at various sites of plasticity throughout the network. This included modification of neuronal participation, burst intensity, excitability, and membrane properties that each showed distinction depending on flexion phase. These results provide a plausible strategy for how these two memories can co-exist in the behavior – there are both shared and unique sites of plasticity to encode learning. To fully appreciate the complex strategies involved in memory overlap, one must observe multiple sites and mechanisms of plasticity at once, or risk mistakenly concluding that the two memories are not both encoded in the same network. It can also not be assumed that all sites of plasticity will utilize the same cellular mechanisms to encode learning.

Invertebrate preparations like *Tritonia,* with their simple behaviors and small nervous systems, allow for easier observation and investigation of multiple sites of plasticity at once. Future investigations interested in memory overlap should consider that even when two memories are encoded within the same network and produce behavioral interference, multiple sites of plasticity may be involved – including sites that are shared and sites that are unique.

## ACKNOWLEDGEMENTS

We would like to acknowledge Margaret Boersma for assisting with behavioral data collection.

## DISCLOSURES

The authors have nothing to disclose.

## FUNDING

NIH R01NS121220 to WNF.

## AUTHOR CONTRIBUTIONS

Conceived and designed research: VKM, ESH, WNF, performed experiments: VKM, ESH, analyzed data: VKM, prepared figures: VKM, drafted manuscript: VKM, edited and revised manuscript: VKM, ESH, WNF, approved final version of manuscript: VKM, ESH, WNF.

## Notes

### Competing Interest Statement

The authors have declared no competing interest.

